# Characterizing and Managing Missing Structured Data in Electronic Health Records

**DOI:** 10.1101/167858

**Authors:** Brett K. Beaulieu-Jones, Daniel R. Lavage, John W. Snyder, Jason H. Moore, Sarah A Pendergrass, Christopher R. Bauer

## Abstract

Missing data is a challenge for all studies; however, this is especially true for electronic health record (EHR) based analyses. Failure to appropriately consider missing data can lead to biased results. Here, we provide detailed procedures for when and how to conduct imputation of EHR data. We demonstrate how the mechanism of missingness can be assessed, evaluate the performance of a variety of imputation methods, and describe some of the most frequent problems that can be encountered. We analyzed clinical lab measures from 602,366 patients in the Geisinger Health System EHR. Using these data, we constructed a representative set of complete cases and assessed the performance of 12 different imputation methods for missing data that was simulated based on 4 mechanisms of missingness. Our results show that several methods including variations of Multivariate Imputation by Chained Equations (MICE) and softImpute consistently imputed missing values with low error; however, only a subset of the MICE methods were suitable for multiple imputation. The analyses described provide an outline of considerations for dealing with missing EHR data, steps that researchers can perform to characterize missingness within their own data, and an evaluation of methods that can be applied to impute clinical data. While the performance of methods may vary between datasets, the process we describe can be generalized to the majority of structured data types that exist in EHRs and all of our methods and code are publicly available.

## BACKGROUND AND SIGNIFICANCE

Missing data present a challenge to researchers in many fields and this challenge is growing as datasets increase in size and scope. This is especially problematic for electronic health records (EHRs), where missing values frequently outnumber observed values, and the absence of an observation can be caused by a variety of mechanisms that may or may not be informative. EHRs were designed to record and improve patient care and streamline billing, and not as resources for research[1] thus there are significant challenges using these data to gain a better understanding of human health. As EHR data become increasingly utilized as a source of phenotypic information for biomedical research[2] it is crucial to develop strategies for coping with missing data.

Clinical laboratory assays are a particularly rich data source within the EHR, but they also tend to have large amounts missing data. These data may be missing for many different reasons. Some tests are used for routine screening but screening may be biased. Other tests are only conducted if they are clinically relevant to very specific ailments. Patients may also receive care at multiple healthcare systems, resulting in information gaps at each institution. Age, sex, socioeconomic status, access to care, and medical conditions can all impact how comprehensive the data is for a given patient.

Aside from the uncertainty associated with a variable that is not observed, many analytical methods, such as regression or principal components analysis are designed to operate on a complete dataset. The easiest way to implement these procedures is to remove variables with missing values or remove individuals with missing values. Eliminating variables is justifiable in many situations, especially if a given variable has a large proportion of missing values, but doing so may restrict the scope and power of a study. Removing individuals with missing data is another option known as complete case analysis. This is generally not recommended unless the fraction of individuals that will be removed is small enough to be considered trivial, or there is good reason to believe that the absence of a value is due to random chance. If there are any differences between individuals with and without observations, complete case analysis will be biased. For example, if only patients with severe symptoms receive a certain test, removing patients with missing values is equivalent to removing the healthy patients.

An alternative is to fill the missing fields with estimates. This process, called imputation, requires a model that makes assumptions about why only some values were observed. These missingness mechanisms fall somewhere in a spectrum between three scenarios (Figure 1). When data is missing in a manner completely unrelated to both the observed and unobserved values, it is considered to be missing completely at random (MCAR) [3,4]. When data are MCAR, the observed data represent a random sample of the population, but this is rarely encountered in practice. Conversely, data missing not at random (MNAR) refers to a situation where the probability of observing a data point depends on the value of that data point [5]. In this case, the mechanism responsible for the missing data is biased and should not be considered ignorable [6]. For example, rheumatoid factor is an antibody detectable in blood, and the concentration of this antibody is correlated with the presence and severity of rheumatoid arthritis. This test is typically performed only on patients with some indication of rheumatoid arthritis. Thus, patients with high rheumatoid factor levels are more likely to have rheumatoid factor measures.

**Figure 1.**
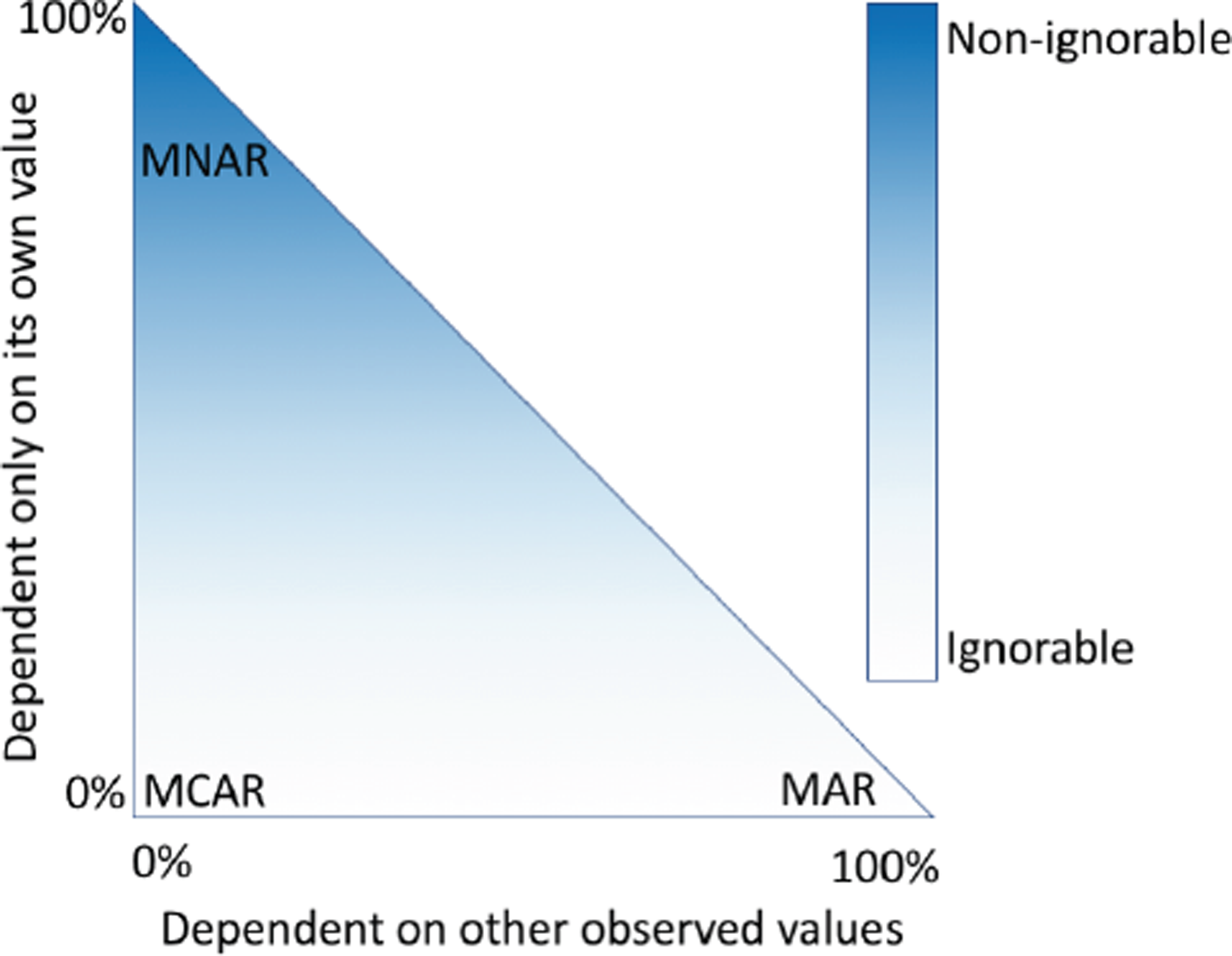
Mechanisms of missing data. Two general paradigms are commonly used to describe missing data. Missing data are considered ignorable if the probability of observing a variable has no relation to the value of the observed variable, and are considered non-ignorable otherwise. The second paradigm divides missingness into three categories: missing completely at random (MCAR: the probability of observing a variable is not dependent on its value or other observed values), missing at random (MAR: the probability of observing a variable is not dependent on its own value after conditioning on other observed variables), and missing not at random (MNAR: the probability of observing a variable is dependent on its value, even after conditioning on other observed variables). The X-axis indicates the extent to which a given value being observed depends upon other the values of other observed variables. The Y-axis indicates the extent to which a given value being observed depends upon its own value.

A more complicated scenario can arise when multiple variables are available. If the probability of observing a data point does not depend on the value of that data point, after conditioning on one or more additional variables, then that data is said to be missing at random (MAR) [5]. For example, a variable, *X*, may be MNAR if considered in isolation. However, if we observe another variable, *Y*, that explains some of the variation in *X* such that after conditioning on *Y*, the probability of observing *X* is no longer related to its own value, *X* is said to be MAR. In this way, *Y* can transform *X* from MNAR to MAR (Figure 1). There is no way to prove that *X* is randomly sampled after conditioning on covariates unless we measure some of the unobserved values, but strong correlations, ability to explain missingness, and domain knowledge may provide evidence of that the data are MAR.

Imputation methods assume specific mechanisms of missingness and assumption violations can lead to bias in the results of downstream analyses that can be difficult to predict [7,8]. Variances of imputed values are often underestimated causing artificially low *p*-values [9]. Additionally, for data MNAR, the observed values have a different distribution than the missing values. To cope with this, a model can be specified to represent the missing data mechanism, but these models can be difficult to evaluate and may have a large impact on results. Great caution should be taken when handling missing data, particularly data that are MNAR. Most imputation methods assume that data are MAR or MCAR, but it is worth reiterating that these are all idealized states and real data invariably fall somewhere in between (Figure 1).

## OBJECTIVE

We extracted median values for 143 clinical lab measures from 602,366 patients from the EHR of Geisinger Health System (GHS) and performed a series of analyses to characterize the mechanisms of missingness for these data. We then narrowed our focus to 28 measures and used sampling to create a complete dataset with no missing values. Next, we simulated missingness in the dataset based on four mechanisms: 1) missing completely at random, 2) missing not at random and 3) missing at random and 4) missing based on patterns observed in real data. Finally, we compared several imputation methods across each of these scenarios. Our results show a wide range in the error and variance estimates associated with different imputation methods and this analysis provides an open source framework that other researchers can follow when dealing with missing data.

## MATERIALS AND METHODS

### Source Code

Source code to reproduce the analyses in this work are provided in our repository (https://github.com/EpistasisLab/imputation) under a permissive open source license. In addition, Continuous Analysis[10] was used to generate Docker images matching the environment of the original analysis, and to create intermediate results and logs. These artifacts are freely available (https://hub.docker.com/r/brettbj/ehr-imputation/ and archive version 10.6084/m9.figshare.5165653).

### EHR data processing

All clinical laboratory assays were mapped to LOINC (Logical Observation Identifiers Names and Codes). We restricted our analysis only to outpatient lab results to minimize the effects of extreme results from inpatient and emergency department data. We used all laboratory results between August 8th, 1996 and March 3rd, 2016. We excluded any codes for which less than 0.5% of patients had a result. The resulting dataset consisted of 669,212 individuals and 143 laboratory assays.

We next removed any lab results that occurred prior to patient’s 18th birthday or after their 90th. In cases where a date of death was present, we also removed any lab results that occurred within one year of death as we found that the frequency of observations often spiked during this period and the values for certain labs were altered for patients near death. For each patient’s clinical lab measures, a median date of observation was then calculated based on all of their remaining lab results. We defined a temporal window of observation by removing any lab results that were recorded more than 5 years from the median date. We then calculated the median result of the remaining labs for each patient. Finally, we calculated the mean BMI of each patient, and dropped any patients whose sex was unknown. As each variable had a different scale and many deviated from normality, we applied Box-Cox and Z-transformations to all variables. The final dataset used for all downstream analyses contained 602,366 patients and 146 variables (age, sex, BMI, and 143 laboratory measures).

### Variable selection

We first ranked the labs by total missingness. At each rank, we calculated the percent of complete cases for the set including all lower ranked labs. We also built a random forest classifier to predict the presence or absence of each variable. Based on these results, in conjunction with domain knowledge, we selected 28 variables that provided a reasonable trade-off between quantity and completeness and were deemed to be largely MAR.

### Predicting the presence of data

For each clinical lab, we used the default scikit-learn [11] random forest classifier, to predict whether a specific value would be present or absent. Each lab measure was converted to a binary label vector based on whether the measure was recorded or not. The values of all other labs, excluding co-members of a panel, were used as the training matrix input to the random forest. This process was repeated for each lab test using 10-fold cross validation. Prediction accuracy was assessed by the area under the receiver operator characteristic (AUC ROC).

### Sampling of complete cases

To generate a set of complete cases that resembled the whole population, we randomly sampled 100,000 patients without replacement. We then matched each of these individuals to the most similar patient who had a value for each of the 28 most common labs by matching sex and finding the minimal Euclidean distance of age and BMI.

### Simulation of missing data

Within the sampled complete cases, we selected the data for removal by four mechanisms:

Simulation 1: Missing Completely at Random (MCAR)

We replaced values with NAN at random. This procedure was repeated 10 times each for 10%, 20%, 30%, 40%, and 50% missingness yielding 50 simulated datasets.

Simulation 2: Missing at Random (MAR)

We selected two columns (*A* and *B*) and a quartile. For the values from column *A* within the quartile, we randomly replaced 50% of the values from column *B* with NAN. The procedure was repeated for each quartile and each lab combination yielding 3024 simulated datasets.

Simulation 3: Missing not at Random (MNAR)

We selected a column and a quartile. When the column’s value was in the quartile we replaced it with NAN 50% of the time. This procedure was repeated for each of the 4 quartiles of each of the 28 labs generating a total 112 total simulated datasets.

Simulation 4: Missingness based on real data observations

From our sampled complete cases dataset, we took each patient and matched them to the nearest neighbor, excluding self matches, in the entire set of observed data based on their sex, age, and BMI. We then replaced any lab value with NAN if it was absent for the matched patient in the original data.

### Imputation of Missing Data

Using our simulated datasets (Simulations 1 - 4) we compared 18 common imputation methods (representative 12 methods are shown in primary figures) from the *fancyimpute* [12] and the Multivariate Imputation by Chained Equations (MICE) [13] packages. A full list of imputation methods and the parameters used for each are in Supplementary Table 1.

## RESULTS

Our first step was to select a subset of the 143 lab measures for which imputation would be a reasonable approach. We began by ranking the clinical lab measures in descending order by the number of patients who had an observed value for that lab. At each rank, we plotted the percent of individuals missing a value for that lab as well as the percent of complete cases when that given lab was joined with all of the labs of lower rank. These plots showed that the best tradeoff between quantity of data and completeness would fall between 20 and 30 variables, Figure 2A.

**Figure 2.**
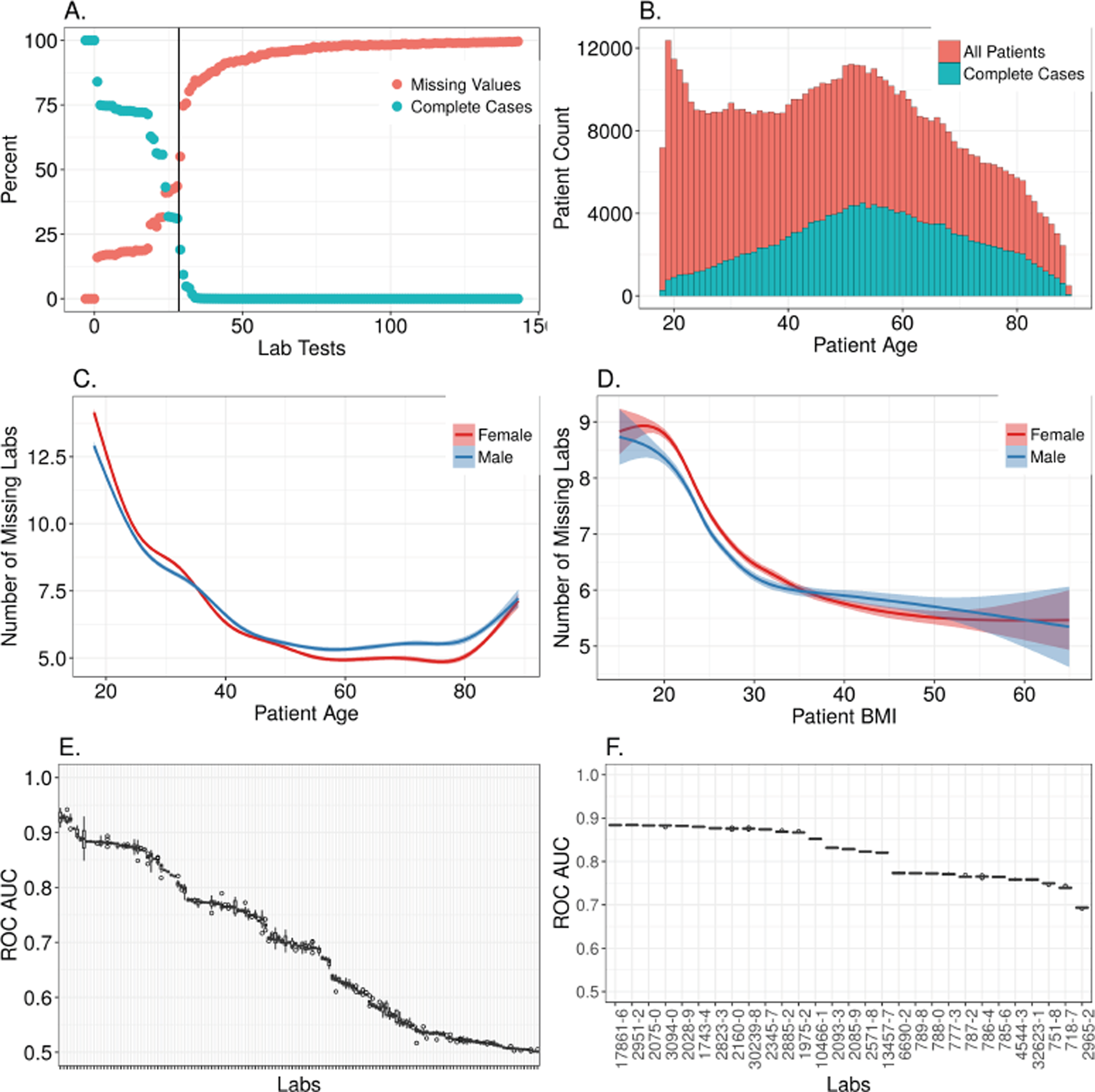
Summary of missing data across 143 clinical lab measures of the Geisinger Health System EHR. **A.)** After ranking the clinical laboratory measures by the number of total results, the percent of patients missing a result for each lab was plotted (red points). At each rank, the percent of complete cases for all labs of equal or lower rank were also plotted (blue points). Only variables with a rank of 75 or less are shown. The vertical bar indicates the 28 labs that were selected for further analysis. **B.)** The full distribution of patient median ages is shown in blue and the fraction of individuals in each age group that had a complete set of observations for labs 1-28 are shown in red. **C.)** Within the 28 labs that were selected for imputation analyses, the mean number of missing labs is depicted as a function of age. **D.)** Within the 28 labs that were selected for imputation, the mean number of missing labs is depicted as a function of BMI. **G.)** Accuracy of a random forest predicting the presence or absence of all 143 laboratory tests. **H.)** Accuracy of a random forest predicting the presence or absence of the top 28 laboratory tests.

**Table 1.** LOINC codes and descriptions of the most frequently ordered clinical laboratory measurements. The assays are ranked from the most common to the least.

| LOINC | Description |
| --- | --- |
| 718-7 | Hemoglobin [Mass/volume] in Blood |
| 4544-3 | Hematocrit [Volume Fraction] of Blood by Automated count |
| 787-2 | Erythrocyte mean corpuscular volume [Entitic volume] by Automated count |
| 786-4 | Erythrocyte mean corpuscular hemoglobin concentration [Mass/volume] by Automated count |
| 785-6 | Erythrocyte mean corpuscular hemoglobin [Entitic mass] by Automated count |
| 6690-2 | Leukocytes [# /volume] in Blood by Automated count |
| 789-8 | Erythrocytes [# /volume] in Blood by Automated count |
| 788-0 | Erythrocyte distribution width [Ratio] by Automated count |
| 32623-1 | Platelet mean volume [Entitic volume] in Blood by Automated count |
| 777-3 | Platelets [# /volume] in Blood by Automated count |
| 2345-7 | Glucose [Mass/volume] in Serum or Plasma |
| 2160-0 | Creatinine [Mass/volume] in Serum or Plasma |
| 2823-3 | Potassium [Moles/volume] in Serum or Plasma |
| 3094-0 | Urea nitrogen [Mass/volume] in Serum or Plasma |
| 2951-2 | Sodium [Moles/volume] in Serum or Plasma |
| 2075-0 | Chloride [Moles/volume] in Serum or Plasma |
| 2028-9 | Carbon dioxide, total [Moles/volume] in Serum or Plasma |
| 17861-6 | Calcium [Mass/volume] in Serum or Plasma |
| 1743-4 | Alanine aminotransferase [Enzymatic activity/volume] in Serum or Plasma by With P-5'-P |
| 30239-8 | Aspartate aminotransferase [Enzymatic activity/volume] in Serum or Plasma by With P-5'-P |
| 1975-2 | Bilirubin.total [Mass/volume] in Serum or Plasma |
| 2885-2 | Protein [Mass/volume] in Serum or Plasma |
| 10466-1 | Anion gap 3 in Serum or Plasma |
| 751-8 | Neutrophils [# /volume] in Blood by Automated count |
| 2093-3 | Cholesterol [Mass/volume] in Serum or Plasma |
| 2571-8 | Triglyceride [Mass/volume] in Serum or Plasma |
| 2085-9 | Cholesterol in HDL [Mass/volume] in Serum or Plasma |
| 13457-7 | Cholesterol in LDL [Mass/volume] in Serum or Plasma by calculation |

As age, sex, and BMI have a considerable impact on what clinical lab measures are collected, we evaluated the relationship between missingness and these covariates (Figure 2 B-D). We also used a random forest approach to predict the presence or absence of each measure based on the values of the other observed measures. MCAR data is not predictable, resulting in ROC AUC near 0.5. Only 38 of the 143 labs had ROC AUCs less than 0.55 (Figure 2E). Very high ROC AUC are most consistent with data that are MAR. For the top 30 candidate labs based on the number of complete cases, the mean ROC AUC was 0.82. This suggested that the observed data could explain much of the mechanism responsible for the missing data within this set. We ultimately decided not to include the 29^th^ ranked lab, specific gravity of urine (2965-2), since it had a ROC AUC of only 0.69 and is typically used for screening only within urology or nephrology departments (R. Levy MD, personal communication). We did include the lipid measures (the 25^th^-28^th^ ranked labs) since they had ROC AUC values near 0.82 and they are recommended for screening of all patients depending on age, sex, and BMI[14]. Our data confirm that age, sex, and BMI are all predictive of the presence of lipid measures (Supplementary Figure 1 A-B).

To assess the accuracy of imputation methods, we required known values to compare with imputed values. Thus, we generated a set of complete cases from the 28 variables we selected based on our characterization of data missingness. Since the set of complete cases differed from the broader population (Figure 2 B-D), we used sampling and K-nearest neighbors matching to generate a sample of the complete cases that resembled the entire population. We then simulated missing data within this set based on 4 mechanisms: MCAR, MAR, MNAR, and realistic patterns based on the original data.

We next evaluated our ability to predict the presence of each value in the simulated datasets so that we could compare patterns with the real data. These simulations confirmed that our MCAR simulation had low ROC AUC (Figure 3A). The MAR data (Figure 3B) and MNAR data (Figure 3C) were often well predicted, particularly for the MAR data, and when data were missing from the tails of distributions. The AUCs rarely exceeded 0.75 in the MNAR simulations while values above 0.75 were typical in the MAR simulations. This provided additional support to our decision to include the top 28 lab measures, since they all had AUCs between 0.9 and 0.75, which was outside the range of nearly all MNAR simulations (Figure 2F and Figure 3C).

**Figure 3.**
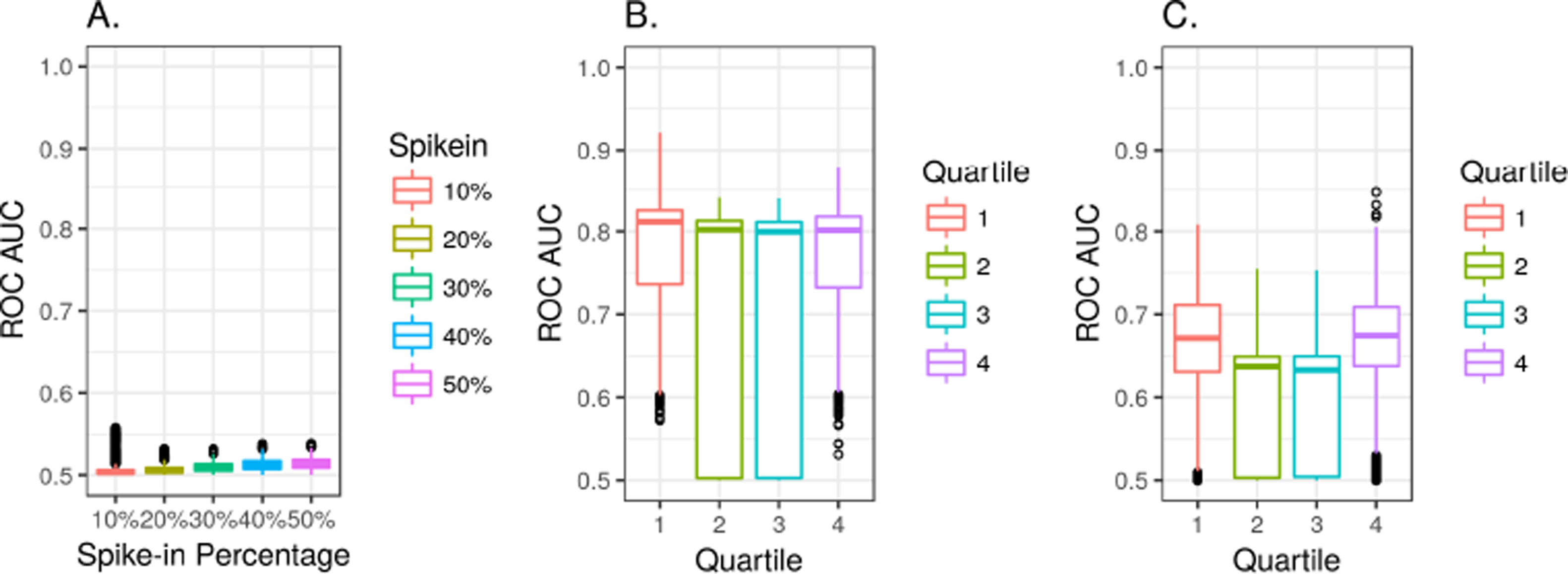
Receiver operating characteristic area under the curve (ROC AUC) of a random forest predicting whether data will be present or missing. **A.)** MCAR simulation **B.)** MAR simulation **C.**) MNAR simulation.

We chose to test the accuracy of several methods from two popular and freely available libraries: the Multivariate Imputation by Chained Equations (MICE) package for R and the *fancyimpute* library for python. We first applied each of these methods across simulations 1-3. For each combination, the overall root mean squared errors are depicted in Figure 4. A breakdown of all the methods and parameters are shown in Supplementary Table 1 with results in Supplementary Figures 3-21.

**Figure 4.**
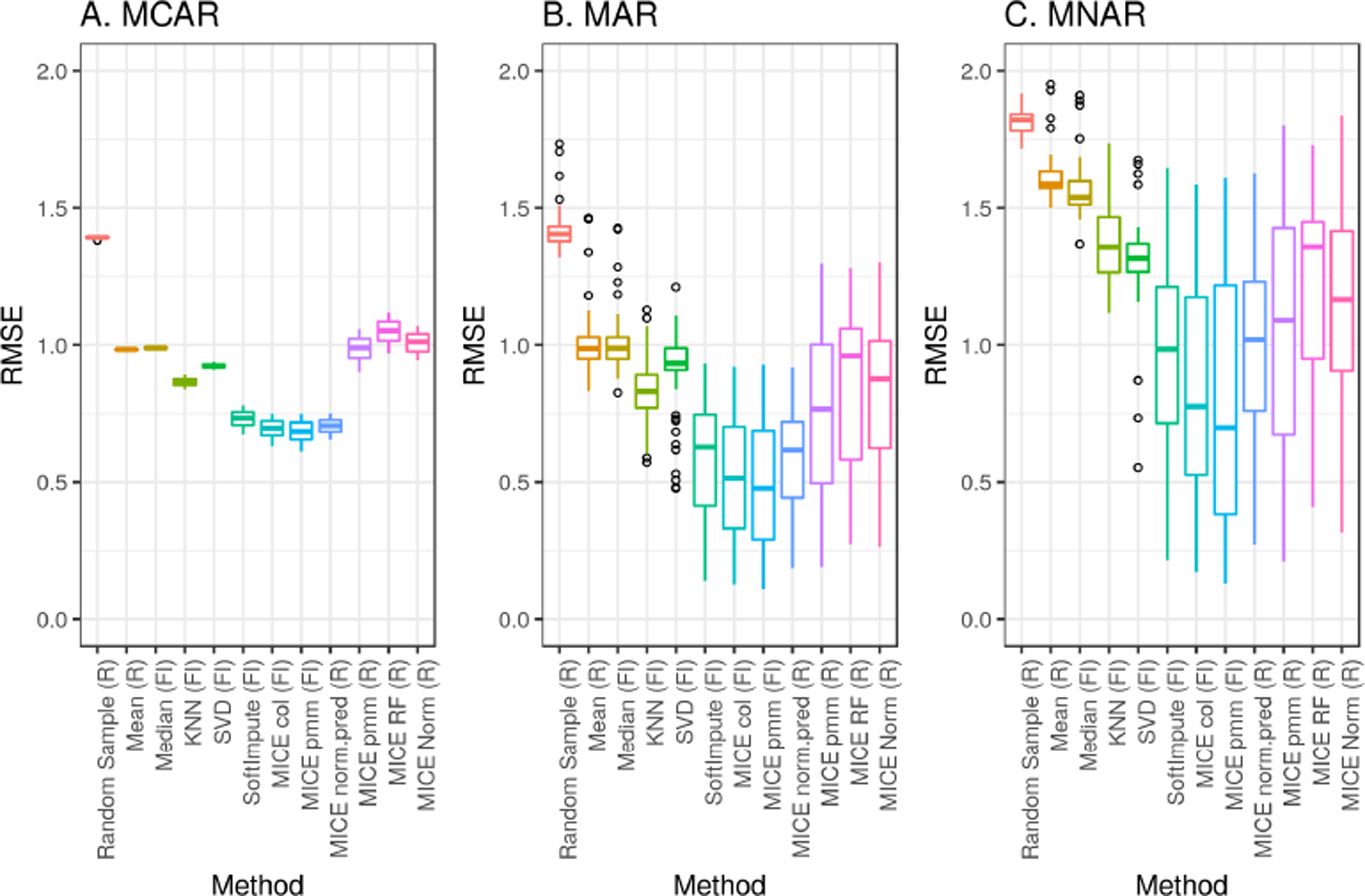
Imputation accuracy measured by RMSE across simulations 1-3. **A.)** Missing Completely at Random (MCAR) **B.)** Missing at Random (MAR) **C.)** Missing Not at Random (MNAR)

We next measured imputation accuracy based on the patterns of missingness that were observed in the real data (Figure 5). The main difference compared to simulations 1-3 was lower error for some of the deterministic methods (Mean, Median, and KNN). It is worth mentioning that the error was highly dependent upon the variable that was being imputed. Specifically, for the fancyimpute MICE PMM method, multicollinearity within some of the variables caused convergence failures that led to extremely large errors (Figure 5, method 8). These factors were relatively easy to address in the R package MICE-PMM method by adjusting the predictor matrix [13].

**Figure 5.**
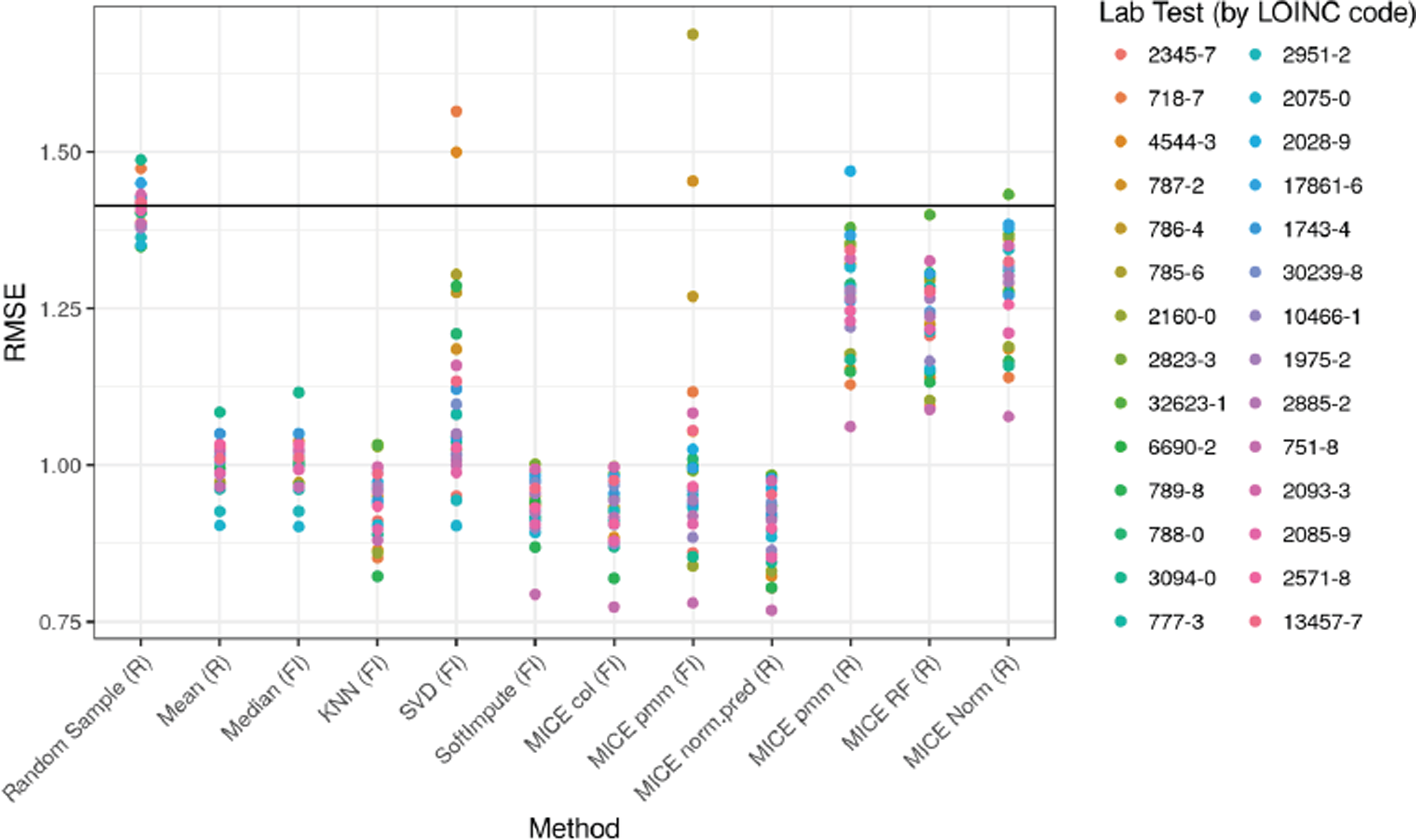
Imputation error (RMSE) for a subset of 10,000 patients from simulation 4. Twelve imputation methods were tested (X-axis) and colors indicate how the error varied between different lab measures (LOINC codes). The black line shows the theoretical error from random sampling.

In addition to evaluating the accuracy of imputation, it is also important to estimate the uncertainty associated with imputation. One approach to address this is multiple imputation, where each data point is imputed multiple times using a nondeterministic method. This allows for the calculation of a confidence interval for any downstream result of interest. To determine if each method properly captured the true uncertainty of the data, we compared the error between an imputed dataset and the observed data with the error between two sets of imputed values for each method (Figure 6). If these errors are equal, then multiple imputation is likely producing good estimates of uncertainty. If, however, the error between two imputed datasets is less than that between each imputed dataset and the known values, then the imputation method is likely underestimating the variance.

**Figure 6.**
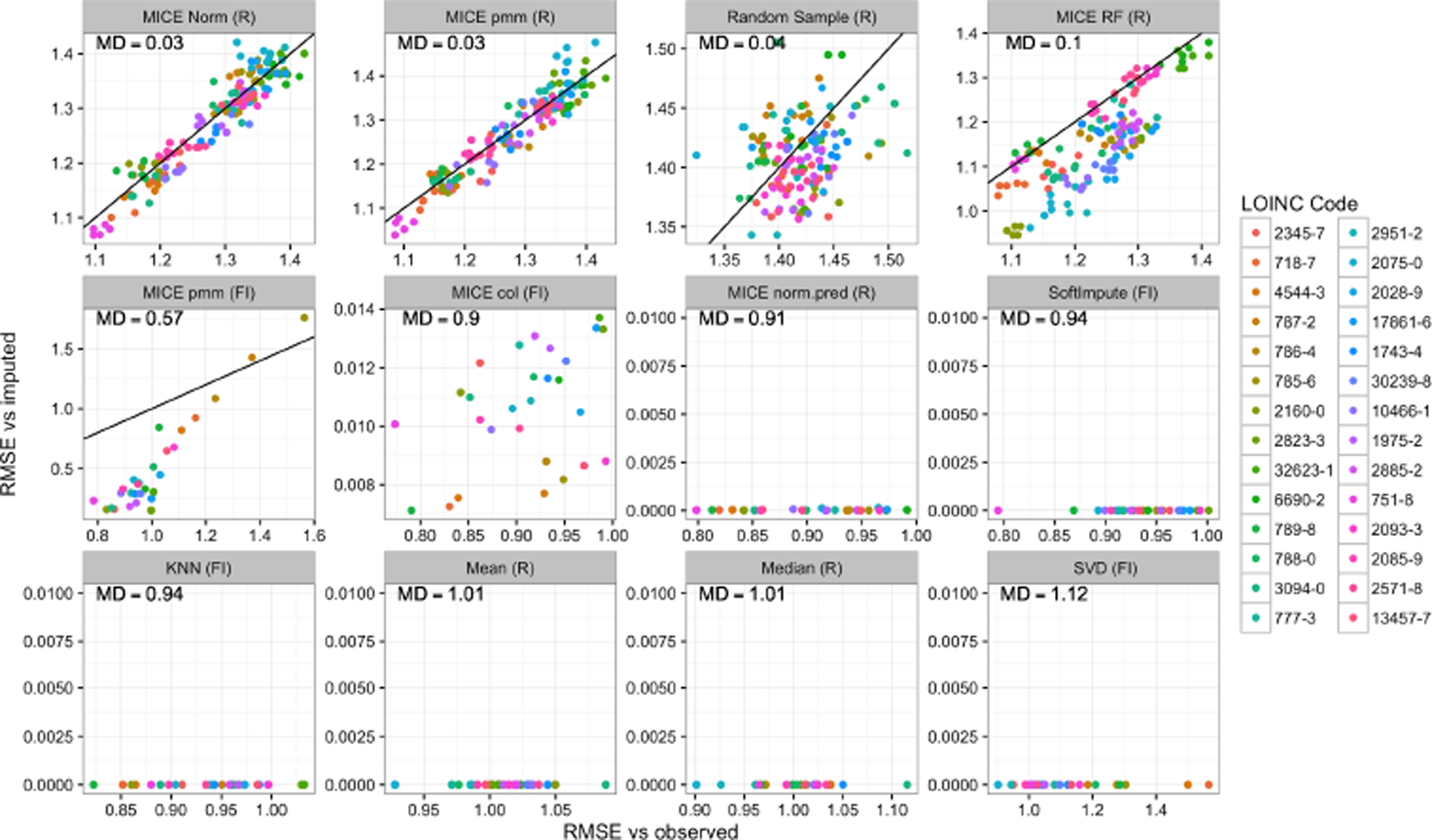
Assessment of multiple imputation for each method. Using simulation 4, we imputed missing values multiple times with each method. The RMSEs between each imputed dataset and the observed values are shown on the X-axis and the RMSEs between two sets of imputed data are shown on the Y-axis. The axis scales vary between panels to better show the range of variation. Each lab test (LOINC) is indicated by the color of the points. The diagonal line represents unity. Panels are ordered by each method’s mean deviation (MD) with unity, indicated in the top left corner of each panel.

Our results (Figure 6) demonstrate that many of the imputation methods are not suitable for multiple imputation. Three of the methods that had the lowest error in the MCAR, MAR, and MNAR simulations (soft impute, MICE col (FI), MICE norm.pred (R)) were found to have minimal variation between imputations. This was also true of KNN, SVD, mean, and median imputation. Only three methods (random sampling, MICE norm (R), and MICE pmm (R)) seemed to have similar error between the multiple imputations and the observed data and thus appear to be unbiased. The latter two had very similar performance and are the best candidates for multiple imputation. Two methods had intermediate performance. MICE RF (R) was similar to several other MICE methods in terms of error relative to the observed data but it produced slightly less variation between each imputed dataset. This seemed to affect some variables more than other but there was no obvious pattern. The MICE pmm (FI) was not deterministic but it did seem to achieve low error at the expense of increased bias. In this case, the variables that could be imputed with the lowest error also seemed to have the most bias. Since this method claims to be a reimplementation of the MICE pmm (R) method, this may be due to multicolinearity among the variables that could not easily be accounted for as there was no simple way to alter the predictor matrix.

## DISCUSSION

It is not possible, or even desirable, to choose “the best” imputation method. There are many considerations that may not be generalizable between different sets of data; however, we can draw some general conclusions about how different methods compare in terms of error, bias, complexity, and difficulty of implementation. Based on our results, there seem to be three broad categories of methods.

The first category is the simple, deterministic, methods. These include mean or median imputation and K-nearest neighbors. The idea behind these methods is that the central tendency of a distribution will be good guess for any unobserved data point. Imputing mean or median values is very easy to implement this but may lead to severe bias and large errors if the unobserved data are more likely to come from the tails of the observed distribution (Figure 3A-C, methods 2-5). This will also cause the variance of the distribution to be underestimated if more than a small fraction of the data is missing. Since these methods are deterministic, they are also not suitable for multiple imputation since no estimate of the uncertainty in the results of any downstream analyses can be made (Figure 6, bottom row).

KNN is similar to mean imputation but based on the idea that there may be groups of individuals that are similar to each other. The value of a missing data point can be estimated by identifying other individuals who have that measure and appear similar to the unmeasured individual based on values of variables observed in both. A group of similar individuals are thus identified and their values are averaged to provide an estimate of the missing value. This generally provides lower error than taking the mean of all individuals, but the choice of K can be difficult to specify. Our simulations suggest that the optimal value can range from less than 1% of the population to more than 50% the population depending on the mechanism of missingness.

KNN is a popular choice for imputation and has been shown to perform very well in some types of data [15,16] but it was not particularly well suited for our data, regardless of the choice of K. This may due to issues of data dimensionality [17] or it may be that human beings do not fall into well separated groups based on their clinical lab results. This method is also not currently suitable for large datasets. The first step is to build a distance matrix for all pairs of individuals that is stored in RAM, and the size of the distance matrix scales with n^2^.

There are also many different methods for calculating distance and the optimal choice may vary widely from one type of data to another. The method that we implemented uses the mean squared linear distance across all pairs of shared observations. While this is probably the best choice when the number of shared features varies between individuals, it assumes that all variables are equal in their ability to capture similarities between individuals regardless of what variable is being imputed. This is certainly not a realistic assumption for our dataset which further points to the fact that imputation is not plug and play and analysis must be done before handling missing data.

The second broad category of algorithms could be called the sophisticated, deterministic methods. These include SVD, soft impute, and MICE col/norm.predict. They tend to rely on either multivariate regression and/or projection of the data into a space of lower dimension. SVD performed poorly compared to its counterparts and sometimes produced errors greater than simple random sampling (Figure 5, method 5). The reasons for this are not clear, but we cannot currently recommend this method. Soft impute and MICE col/norm.predict were among the lowest error methods in all of our simulations (Figure 5, methods 6-7). The main limitation of these methods is that they cannot be used for multiple imputation (Figure 6, middle row).

The third broad category of algorithms were the stochastic methods which included random sampling and most of the remaining methods in the MICE library. The random sampling method almost always produced the highest error (Figure 4-5, method 1) but it has the advantage of being easy to implement and it requires no parameter selection. The MICE methods based on predictive mean matching, random forests, and Bayesian linear regression tended to perform similarly in terms of error in most of our simulations (Figure 4-5, methods 10-12).

Imputation methods that involve some type of stochastic sampling allow for a fundamentally different type of analysis called multiple imputation. In this paradigm, multiple imputed datasets (a minimum of 3 and often 10-20 depending of the percentage of missing data)[18–20] are generated and each is analyzed in the same way. At the end of all downstream analyses, the results are then compared. Typically, the ultimate result of interest is supported by a p-value, a regression coefficient, an odds ratio, etc. In the case of a multiply imputed dataset, the researcher will have several test statistics that can be used to estimate a confidence interval for the result.

Multiple imputation has been gaining traction over the years and the MICE (multiple imputation by chained equations) package has become one of the most popular choices for implementing this procedure. This package is very powerful and very well documented[13] but like all methods for imputation, caution must be exercised. There also seems to be some confusion surrounding the concept of MICE. It is not a single algorithm but rather a framework for applying a variety of algorithms. Each missing value for a variable of interest is imputed by considering the other variables that were observed for that individual, the observed values of the variable of interest in other individuals, and/or the relationships between the variables. This procedure is applied for each missing value in one variable, and then to each subsequent variable. This entire process is then repeated for a number of iterations such that the values imputed in the first iteration can update the estimates for the second iteration. The result is a chain of imputed datasets and this entire process is typically performed in parallel so that multiple chains are generated.

In MICE, there are a number of choices that must be made and care should be taken to evaluate the results. The first obvious choice is the method (i.e. equation). Many methods are available in the base package, additional methods can be added from other packages [https://cran.r-project.org/web/packages/miceadds/index.html], and users can even define their own. These methods could be extended in theory to include any of the previously described algorithms and the base package already includes random sampling and mean imputation. We thoroughly evaluated three methods in the context of our dataset: predictive mean matching (pmm), bayesian linear regression (norm), and random forest (rf).

PMM is the default choice and popular since it can be used on a mixture of numeric and categorical variables. We found PMM to have a good trade-off between error and bias, but for our dataset it was critical to remove several variables from the predictor matrix due to strong correlations (R > 0.85) and multicolinearity. Bayesian regression performed similarly but was less sensitive to these issues. If a dataset contains only numeric values, Bayesian regression may be a safer option. Random forest tended to produce results that were slightly biased for a subset of the variables without an appreciable reduction in error. Aside from random sampling, none of the other methods we evaluated were suitable for multiple imputation (Figure 6).

## CONCLUSION

There are many factors that must be considered when analyzing a dataset with missing values. This starts by determining whether each variable should be considered at all. Two good reasons to reject a variable are if has too many missing values or if it is likely to be MNAR. There is no simple rule to determine the amount of missingness that can be tolerated but it is desirable to select a set of variables such that there are enough complete cases to get a reasonable estimate of how each variable relates to every other variable. While it is impossible to completely rule out the possibility that a variable is largely MNAR, it is possible to model the presence or absence of values as a function of the observed values. If such a model has good predictive power, this provides some evidence that the variable may be MAR and the assumptions of most imputation methods may not be severely violated. If this procedure fails, then the data likely lie somewhere in the spectrum between MCAR and MNAR. The former case also meets the assumptions of most imputation methods but tends to be rare in practice. If a variable is MNAR, it may still be possible to impute, but the mechanism of missingness should be explicitly modeled and a sensitivity analysis is recommended to assess how much impact this could have on the final results[21,22]. While a statistical model of the mechanism of missingness is useful, there is no substitute for a deep familiarity with data at hand and how it was generated.

Once the data are selected, the main decision is how to impute. Many methods have been proposed and each has limitations. Depending upon the number of individuals, the relationships between the individuals, the number of variables, the relationships among the variables, the patterns of missingness, and the mechanisms of missingness, the performance of any given method may vary dramatically. In order to assess the performance an imputation method it should ideally be tested in a realistic setting. Great care should be taken to construct a set of complete data that closely resemble all of the relevant characteristics of the data that one wishes to impute. Similar care should then be taken to remove some of this data in ways that closely resemble the observed patterns of missingness. If this is not feasible, it may be possible to simulate a variety of datasets representing a range of possible data structures and missingness mechanisms. Any available imputation methods can then be applied to the simulated data and error between the imputed data and their known values provide a metric of performance.

While the minimization of error in the imputed data is the primary goal, a singular focus on this objective is likely to lead to bias. For each missing value, it is also important to estimate the uncertainty associated with it. This can be achieved by multiple imputation using an algorithm that incorporates stochastic processes. Multiple imputation has become the field standard because it provides confidence intervals for the results of downstream analyses. One should not naively assume that any stochastic process is free of bias. It is important to check that multiple imputation is providing variability that corresponds to the actual uncertainty of the imputed values using a set of simulated data.

## Acknowledgments

We thank Casey S. Greene (University of Pennsylvania) for his helpful discussions.

## Funding

This work was supported by the Commonwealth Universal Research Enhancement (CURE) Program grant from the Pennsylvania Department of Health. B.K.B.-J. and J.H.M. were also supported by US National Institutes of Health grants AI116794 and LM010098 to J.H.M.

## Author Contributions

B.K.B.-J., J.H.M., S.A.P. and C.R.B. and conceived of the study. D.L. and J.S. performed data processing. B.K.B.-J. and C.R.B. performed analyses. B.K.B.-J., S.A.P. and C.R.B. wrote the manuscript and all authors revised and approved the final manuscript.

## Competing Interests

The authors have no competing interests to disclose.

## Source code availability

All source code is available via github (https://github.com/epistasislab/imputation).

